# SAPH-ire TFx – A Recommendation-based Machine Learning Model Captures a Broad Feature Landscape Underlying Functional Post-Translational Modifications

**DOI:** 10.1101/731026

**Authors:** Nolan English, Matthew Torres

## Abstract

Protein post-translational modifications (PTMs) are a rapidly expanding feature class of significant importance in cell biology. Due to a high burden of experimental proof, the number of functional PTMs in the eukaryotic proteome is currently underestimated. Furthermore, not all PTMs are functionally equivalent. Therefore, computational approaches that can confidently recommend the functional potential of experimental PTMs are essential. To address this challenge, we developed *SAPH-ire TFx* (https://saphire.biosci.gatech.edu/): a multi-feature neural network model and web resource optimized for recommending experimental PTMs with high potential for biological impact. The model is rigorously benchmarked against independent datasets and alternative models, exhibiting unmatched performance in the recall of known functional PTM sites and the recommendation of PTMs that were later confirmed experimentally. An analysis of feature contributions to model outcome provides further insight on the need for multiple rather than single features to capture the breadth of functional data in the public domain.

**Supplementary Information:** See Tables S1-S6 & Figures S1-S4.

## INTRODUCTION

Post-translational modifications (PTMs), chemical or proteinaceous alterations to amino acid residues in a protein, have the potential to expand the function and regulatory control of proteins beyond the limits of the genome (Prabakaran et al., 2012). PTMs can act on long or short timescales that allow for dynamic control and response of a cellular proteome to changing environments or cellular phases that ultimately shape cellular phenotype, often by modulating changes in protein interaction, localization, or stability (Csizmok and Forman-Kay, 2018). Concomitantly, disruption to either the amino acid or modification of highly functional PTM sites can contribute to cellular dysfunction and disease (Gibson et al., 2010; Reimand et al., 2015; Reimand and Bader, 2014).

The scientific community has witnessed an exponential increase in PTM data over the last 15 years, fueled by high-throughput mass spectrometry that has identified hundreds of different PTM types occurring on nearly all of the 20 common amino acids. However, the rate at which PTM data is generated – a parallel process involving hundreds of independent labs – far surpasses the rate at which it is being curated and/or processed for interpretation – a task undertaken by a much smaller set of labs and institutions (Chen et al., 2017; Pascovici et al., 2018). A longstanding question emerging from these efforts is whether all PTMs (*detected accurately*) are functionally important – a question not easily answered due to the high burden of experimental evidence needed to prove functionality, which involves significant time, cost, and specific expertise for any given protein. These challenges are compounded by unnecessary redundancy in experimental effort and the tendency of most labs not to report non-functional results. Although not as commonly addressed in the literature, lack of PTM-centric user-friendly visualization and organization tools – with or without computational enhancements – also raises significant barriers to PTM data accessibility and interpretation. These underlying challenges limit the view of what are an are not likely important modifications and this tends to promote a perspective that the study of PTMs is risky and quite possibly not worth the effort.

Computational approaches aimed at the functional prioritization, or rank-based sorting, of PTMs using single PTM site features have made a tangible impact on the discovery of several new regulatory elements in proteins. Indeed, functional significance of PTM sites that are evolutionarily conserved – especially across a great phylogenetic distance – have proven to be more likely functional for the protein families in which they are found (Beltrao et al., 2012; Landry et al., 2009; Strumillo et al., 2019). Similarly, co-localization was shown to be predictive for co-regulatory phosphorylation-dependent ubiquitination (Minguez et al., 2015, 2013, 2012). Lastly, protein structural features such as solvent accessibility or PTM proximity to catalytic residues has proven to be a useful filter for functional modifications (Dewhurst et al., 2015; Johnson et al., 2015). Despite these successes, not all PTMs with experimental evidence of function are highly conserved, co-localized, or are near catalytic pockets or other important protein structures. Indeed, cases wherein a PTM’s potential for function is easily predicted by one of these co-occurring features alone may be considered “low-hanging fruit”.

Machine learning models that incorporate multiple PTM site features have shown promise in capturing a larger proportion of the functional PTM population in eukaryotes (Ochoa et al., 2020; Torres et al., 2016; Xiao et al., 2016), and have enabled the identification of functional PTMs not readily identifiable through single feature analyses alone (Dewhurst and Torres, 2017). However, most models have limited potential to inform the broad range of functionality likely to exist across the Eukaryotic kingdom as most exclude all but one type of PTM – usually phosphorylation – despite the ample evidence of many other regulatory modifications and sites. Existing models are also rank-based, in which model output places PTMs in a competitive hierarchy of functional importance. Within these models, PTMs for which functional evidence already exists end up being broadly distributed in the scoring regime and with only a small fraction of candidates rising to the top. This severely limits the utility of rank-based methods for identifying PTMs of putative function as only the most extreme outliers can be confidently chosen.

We hypothesize that capturing the breadth of function that exists naturally in biology can benefit from the use of *inclusive* models that incorporate data from many different PTM types, PTM site features, and functional consequences. Here we test this hypothesis through the development, characterization and application of a new machine learning model, SAPH-ire TFx, and a complimentary interactive web-based resource and API (https://saphire.biosci.gatech.edu) to enable PTM data visualization. The model is recommendation rather than rank based and has been applied to 512,015 total unique PTMs of which ∼12,000 have been validated as functional *a priori* across 763 eukaryotic organisms. Extension of the results to experimental PTMs of unknown function suggest that as many as half of them do not exhibit characteristics of functional PTMs in the public domain.

## RESULTS

### SAPH-ire TFx exhibits robust performance on unexposed datasets

A detailed description of the SAPH-ire TFx model design and architecture is described within materials and methods. Conceptually, the model utilizes feature data extracted from multiple sequence alignment positions that harbor evidence of PTM (called Modified Alignment Positions or *MAPs*), uses these features as inputs into a neural network trained to recognize MAPs harboring functional PTMs (called *known functional MAPs*) (**Figure 1**). For this study, the model was developed on an initial dataset compiled in late 2018, consisting of 435,750 total PTMs of which 9,151 were known functional (the *training dataset*; see materials and methods). We then evaluated its performance on an expanded PTM dataset in which 102,475 unexposed PTMs (3,233 known functional) were added to the original set (i.e. the expanded data was not part of the training nor validation processes employed during model development) (**Figure 2A**).

**Figure 1.**
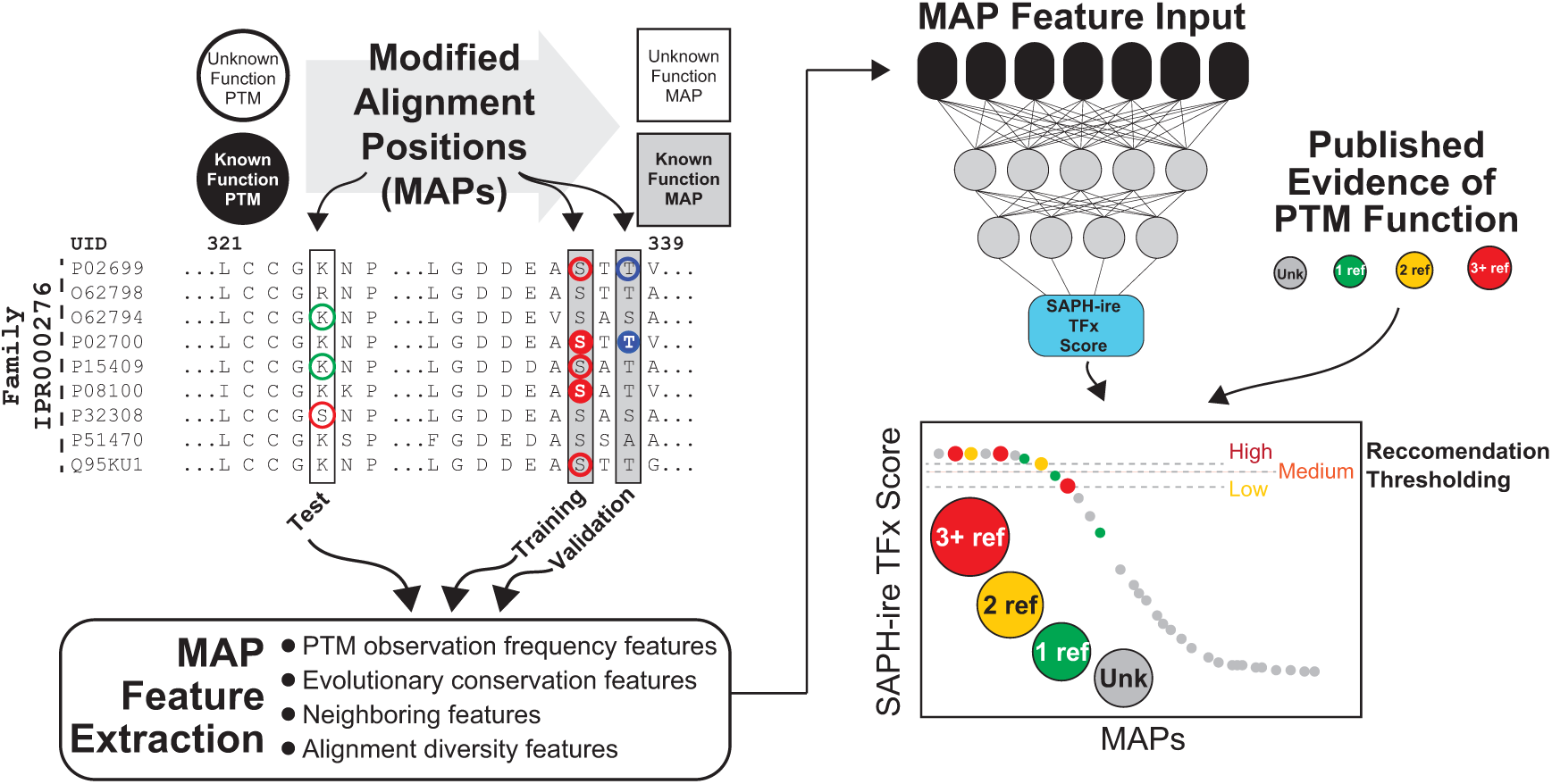
Schematic diagram of the SAPH-ire TFx methodology. PTMs of both unknown and known functional consequence (as determined through curated public record) are organized by full length protein family multiple sequence alignment, creating Modified Alignment Positions (MAPs) of unknown or known function. Known function MAPs are used either for model training and/or model validation (via calculation of model recall) while unknown function MAPs represent the test cases for which a functional impact is not currently known for any of the aligned PTMs. Features are extracted from MAP data and then these features used as inputs into a neural network trained to identify known functional MAPs. At this point the model is blind to whether a MAP is known or unknown. Each MAP (both known and unknown) passes through the model to receive a SAPH-ire TFx output score that ranges from 0 to 1, where 1 indicates a MAP that closely resembles a known functional MAP. After scoring, the status of each MAP as known or unknown function and the sum of literature sources supporting evidence of function (i.e. the Known Function Source Count) is revealed. Model performance is graded and recommendation thresholds generated using recall of known function MAPs as a guide.

**Figure 2.**
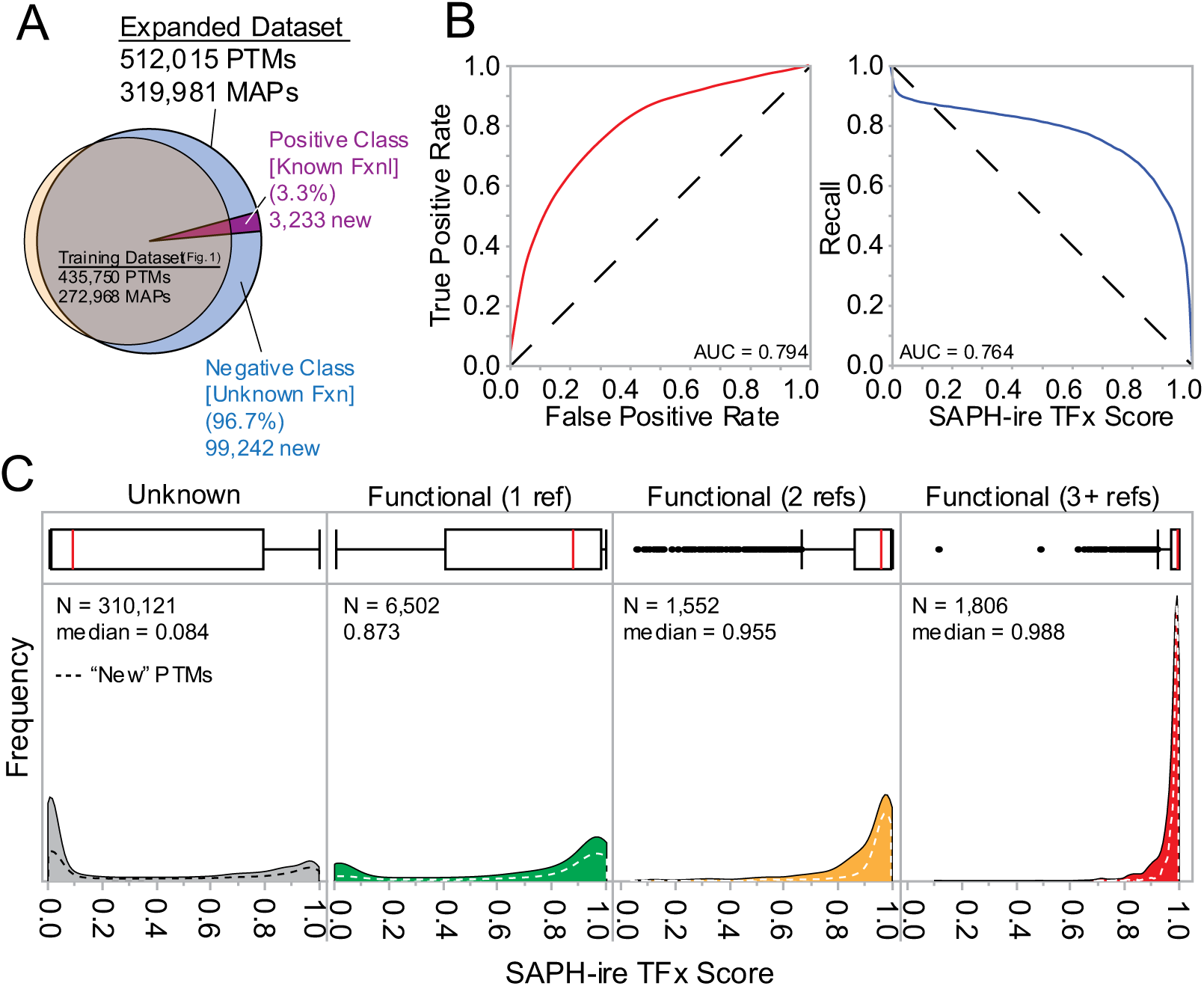
SAPH-re TFx performance on an unexposed dataset. (A) Venn diagram showing the relationship between the training and expanded datasets. The expanded dataset contained 102,475 newly curated PTMs. (B) ROC and recall curves for SAPH-ire TFx results from the expanded dataset. (C) Frequency distribution of SAPH-ire TFx scores relative to true positive status in terms of known function source count (KFSC = 0, 1, 2, or 3+ references). Area contained by solid lines corresponds to the total expanded PTM dataset. Area contained by dashed lines corresponds to model output for unexposed PTMs not contained in the original training dataset. All statistical data shown is aggregated at the MAP level.

To evaluate performance, we used area under the receiver operating characteristic curve (ROC AUC), which reports on model accuracy as well as the recall of known functional MAPs (i.e. true positive data). Overall, model performance on the expanded dataset was better than on the original training dataset for both metrics (AUC_ROC_ 0.794 and AUC_Recall_ 0.764) (*see materials and methods*), suggesting that the addition of new data did not diminish performance (**Figure 2B**). Next, we evaluated the model outcome score distributions for unknown and known functional MAPs. MAPs were first binned by known function source count (*KFSC*) – a count of the unique literature sources containing evidence of functional impact for a PTM within the MAP (not included as a feature in the model). This type of performance evaluation is unique and serves as a proxy for confidence in model output, which should prioritize MAPs that were established as functional *a priori*. The model functioned as intended, showing increasing enrichment of known functional MAPs with increasing model score (**Figure 2C**). Moreover, we observed a decrease in the variance of the prediction with increasing KFSC. These same trends were also evident for the 3,233 unexposed known functional PTMs in the expanded dataset to which the model was unexposed during development, demonstrating the robustness of the model (**Figure 2C**, dashed lines). Taken together, the data show that SAPH-ire TFx is a robust and effective model capable of distinguishing functional PTMs across independent datasets.

### Analysis of feature contributions in the SAPH-ire TFx model: No single feature can capture all known function PTMs

SAPH-ire TFx incorporates 11 features derived from both empirically and biologically relevant features. To understand how the model balances these features to reach its conclusions and to determine if it is overly reliant on any single feature, we sampled 29,859 MAPs and conducted a Linear Interpretability Model Explanation (LIME) analysis to calculate feature contributions for each MAP. We then clustered the samples in this feature space using normal mixtures. Clusters 1 and 3 have an overrepresentation of known functional MAPs within them (70%+) whilst making up less than 9% of the sampled MAPs. In contrast, cluster 2 represents 92% of sampled MAPs but also has a minority population of known functional MAPs (**Figure 3A**). We used principle components analysis (PCA) to understand the differences in feature contributions between each cluster, which can give insight into the how SAPHire-TFx decides its recommendations (**Figure 3B**). We found that 55% of the variance within the sampled MAPs can be explained by PC1 and PC2, with the other 9 principal components contributing marginally to the remaining 45%. Furthermore, we found that cluster 1 has a high variance in terms of both PC1 and PC2, cluster 2 has a low variance in terms of both PC1 and PC2, while cluster 3 is driven mostly by PC1. In depth analysis of the eigenvector values in PC1, reveal the largest contributor is OBSrc, which corresponds to raw observation frequency of PTMs within a MAP, although the count of organisms contributing a residue to the alignment position (AP_OIDc), the count of modified residues in the alignment position (MRc), and the sum of MAPs observed within +/- 7 alignment positions (NC_7) contribute nearly as much (**Figure 3C**). For PC2, the largest contributor is observation source count normalized to the number of members in the family (OBSrc_pf), but is closely followed by modified residue count in neighboring alignment positions (NMRc_7) and disorder tendency (Dis).

**Figure 3.**
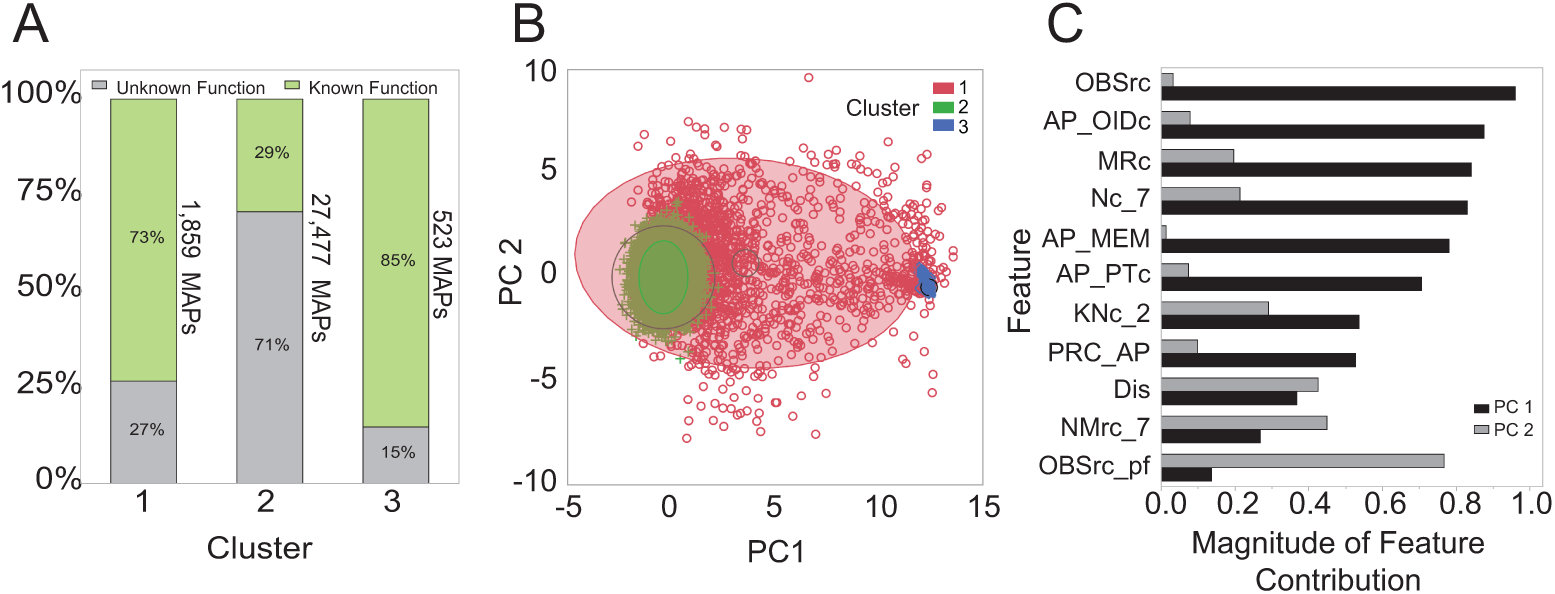
Exploring SAPH-ire TFx’s interpretation of feature space. Normal mixtures clustering of LIME analysis data from 29,859 representative MAPs. (A) Percentage of MAPs within normal mixture clusters that are known to contain a functional PTM. (B) Clusters projected onto two of their principal components. (C) Magnitude of each feature within the principal components shown in B. OBSrc, observation source count; AP_OIDc, alignment position organism ID count; MRc, modified residue count; Nc_7, Neighbor count within +/- 7 alignment positions; AP_MEM, alignment position membership count; AP_PTc, alignment position PTM type count; KNc_2, known functional neighbor count within +/- 2 alignment positions; PRC_AP, PTM residue conservation for the alignment position; Dis, disorder prediction value; NMrc_7, Neighboring modified residue count +/- 7 positions out; OBSrc_pf, observation source count relative to the protein family membership. (*please see detailed feature descriptions in Table S4*)

Extrapolating these characteristics of each principle component reveal that cluster 3, which is highly reliant on PC1, is largely composed of PTMs that are observed frequently globally (OBSrc) and have some weak evidence of functionality from a combination of: one, their proximity to PTMs in neighboring alignment positions; two, the diversity of proteins contributing modified residues to the alignment position; and three, the conservation of the modification across species, for example. Restated, members of this cluster correspond with PTMs that are readily detectable by their detection frequency. Cluster 1 contains PTMs that are observed at a range of frequencies relative to their family and also have strong supporting evidence from other features. Conversely, cluster 2 has little evidence of functionality from the major features of PC1 and PC2, and therefore relies on weak contributions from several features. These results support two major conclusions: one, that single features alone are incapable of capturing the breadth of variation observed for functional PTMs; and two, that SAPH-ire TFx can recognize functional modifications despite this variation.

### PTM-agnostic recommendation thresholding suggests that most PTM sites are not like those we have found are functional thus far

Without further treatment, the SAPH-ire TFx model would be interpreted as a rank-based model and, as described earlier, interpretation of such models is difficult. Implementing recommendation thresholds can be useful to improve interpretation of a model, but simultaneously create boundaries that, if inappropriately placed, can lead to inaccurate predictions. To address these problems, we modeled the tradeoff between true and false positive rates using ROC curves. Our goal was to set a minimum threshold score over which MAPs could be considered having a high chance of functionality. Due to our desire for SAPH-ire scores to be agnostic across different PTM types, we first considered that the selected thresholds must not create a bias in distribution of PTMs occurring above that threshold. To evaluate this, we plotted the percent representation of each of the most common PTMs in the dataset relative to SAPH-ire TFx score (**Figure 4A**). The relative representation of each PTM type deflected significantly above a score of 0.9897 but was stable below this point and above 0.945, the range between which we defined as ideal for thresholding.

**Figure 4.**
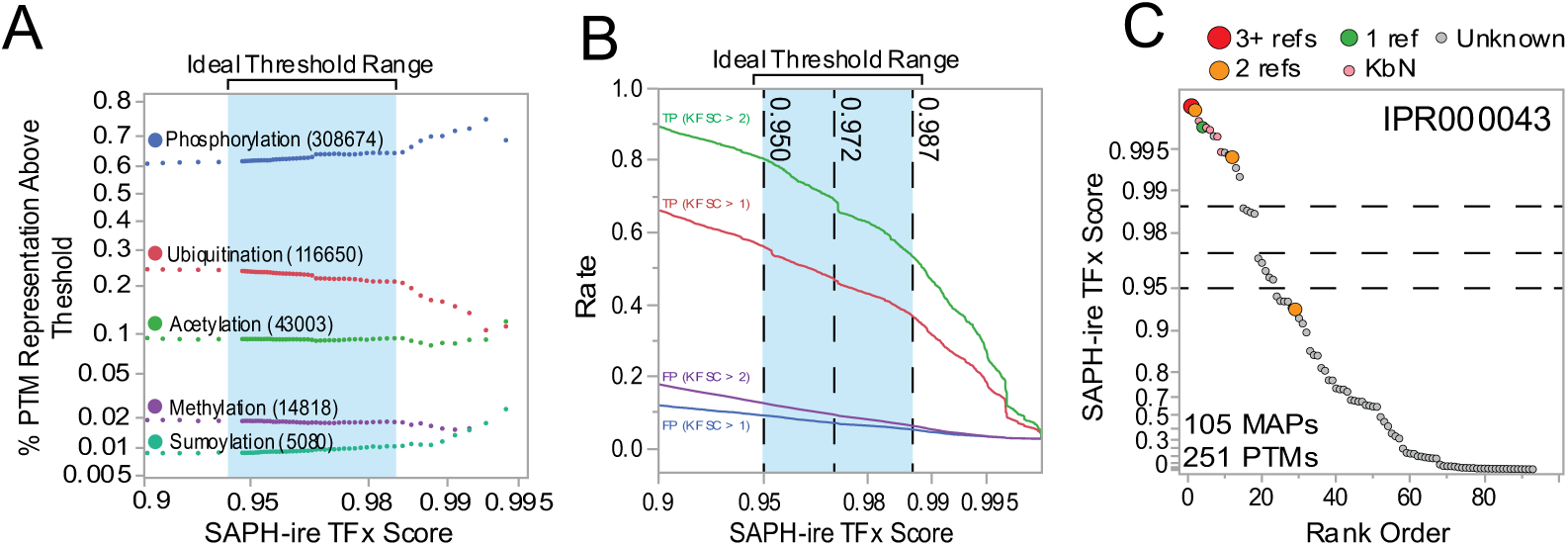
Derivation of SAPH-ire TFx recommendation thresholds. (A) Plot of the percent representation for different PTM types relative to SAPH-ire TFx score, revealing an ideal threshold range inside which no one PTM becomes over or underrepresented. (B) Unfurled ROC curves showing true positive (TP) and false positive (FP) rates above given SAPH-ire score. Rates shown are KFSC > 1 (lower confidence) or KFSC > 2 (higher confidence). Dashed vertical lines represent chosen thresholds where TP and FP rates are as follows (KFSC >2): 0.95 – TP=0.82, FP=0.11; 0.972 – TP=0.7, FP=0.07; 0.987 – TP=0.53, FP=0.04. (C) Representative rank-ordered SAPH-ire TFx plot for family IPR000043 (Adenosylhomocysteinase-like family) with indicated thresholds shown for reference. Shown on an exponential scale to emphasize differences across the scale.

Next, we evaluated the ROC curves for the highest confidence true positive MAPs (KFSC >1, >2) (**Figure S1**), and unfurled each curve to reveal the independent rates for true and false positives with respect to the SAPH-ire TFx score. From these curves, three thresholds were chosen within the ideal range (0.95, 0.9719, and 0.987) that strike a balance between true positive hits and false positive recommendations (**Figure 4B**). These recommendation thresholds provide useful landmarks to interpret SAPH-ire TFx scores for a protein or family of interest, as shown here for family IPR000043 (**Figure 4C**). These thresholds also allow for the evaluation of SAPH-ire in context of other models.

### Benchmarking

SAPH-ire TFx is one of a small number of published algorithms aimed at functional prioritization of PTMs, and the first recommendation-based model for functional PTMs. We therefore sought to draw comparisons with these models to gauge overall performance improvements. Two predominant models currently exist in the public domain: SAPH-ire FPx (S-FPx) (Dewhurst and Torres, 2017) – an 8-feature neural network PTM ranking model; and a Phosphosite Functional Score (PFS) model (Ochoa et al., 2019) – a 59-feature gradient boosting machine learning model trained to identify functional phosphosites.

To evaluate the three models equivalently, we compared model scores for phosphosites represented in all three datasets. PFS was built using a dataset containing 116,268 phosphosites that resulted from selective re-analysis of raw mass spectrometry data files collected from a broad range of eukaryotic organisms (Ochoa et al., 2019). Comparing our source database to the PFS dataset revealed 71% overlap (82,279 phosphosites), however, this number dropped in response to strict protein family membership criteria (*see materials and methods*). Specifically, of PTMs that fall within InterPro whole sequence families, 236,982 represent unique phosphosites that were analyzed by SAPH-ire TFx, and 49,935 of these overlap with ∼43% of the PFS dataset (**Figure 5A**). Inclusion of S-FPx data, which was based on PTMs curated in early 2017, resulted in a final comparable dataset of 24,695 phosphosites.

**Figure 5.**
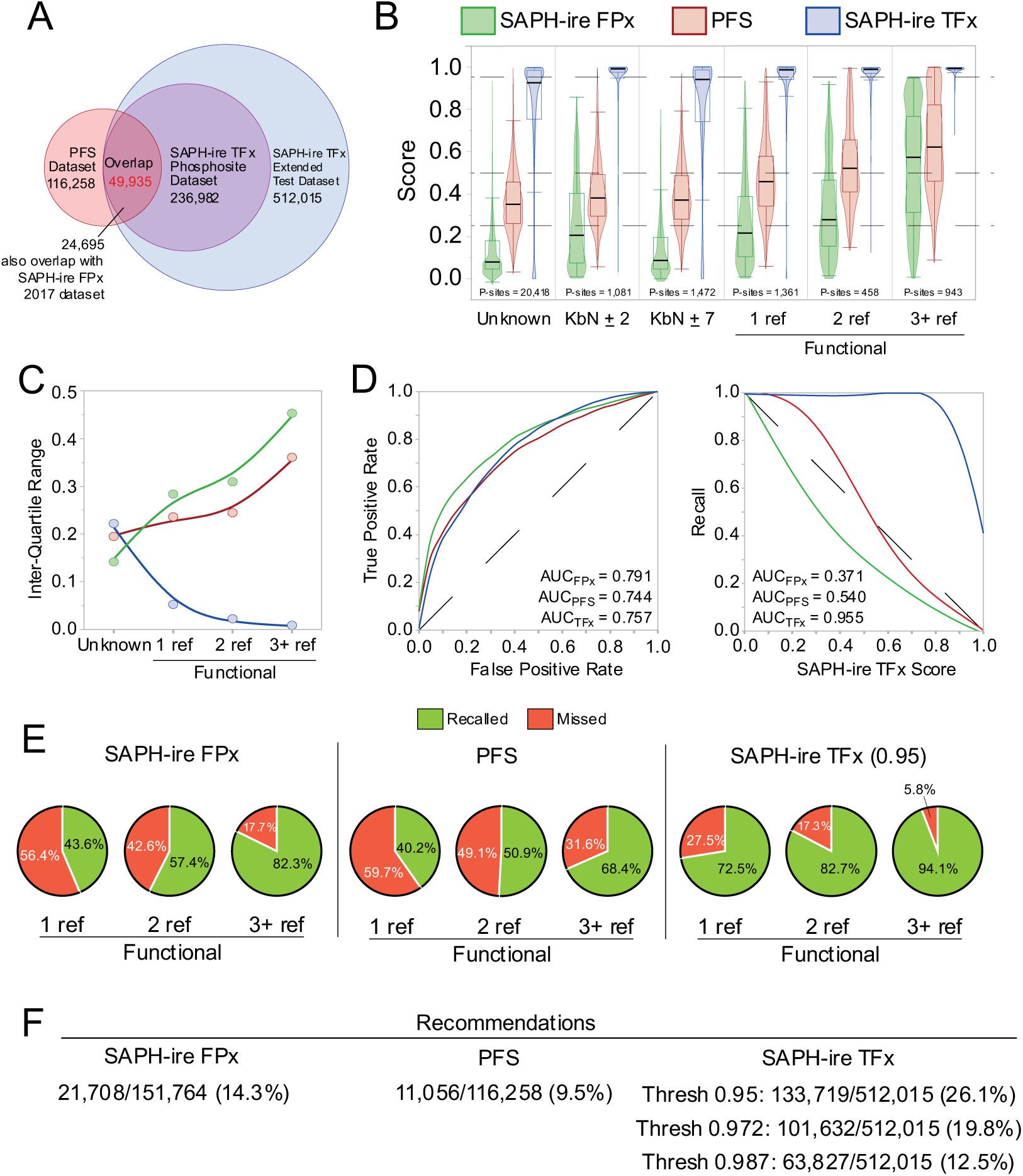
Benchmarking SAPH-ire TFx against existing PTM functional prioritization models. SAPH-ire TFx was compared head-to-head with the two prior machine learning models for functional prioritization: SAPH-ire FPx (Dewhurst and Torres, 2017) and Phosphosite Functional Score (PFS) (Ochoa et al., 2019). (A) Venn diagram describing the overlap between the expanded dataset (reported here) and phosphosite datasets for the other two models. Three-way model comparisons were conducted with 24,695 phosphosites. Pairwise model comparisons (PFS vs. SAPH-ire TFx) were also conducted with 49,935 overlapping phosphosites (Figure S2). (B) Comparison of the score distributions for PTMs binned by category of unknown function, known function (1, 2, or 3+ sources), or known by neighbor (KbN) determined by SAPH-ire TFx protein family alignments. Dashed lines indicate the thresholds quantitatively determined for SAPH-ire TFx or loosely recommended by other models. (C) Inter-quartile range relative to known functional status, based on the distributions shown in B. (D) Comparison of ROC and recall curves for each model. (E) Pie chart representation of the percentage of recalled versus mis-called (Missed) PTMs based on thresholds shown in B [0.95 threshold used for SAPH-ire TFx] (top). Number of recommendations deduced from these percentages applied to the whole dataset for each model (bottom). Recommendations are also shown for each of the thresholds established for SAPH-ire TFx in figure 3.

In general, S-FPx and PFS perform similarly in most respects – in part because they were both rank based models built to maximize ROC AUC but not recall. Both models result in broad and overlapping score distributions that are significantly different but modestly distinct between sites of known and unknown function (**Figure 5B**). This results from broad score distributions that change marginally across bins of increasing KFSC. S-FPx tends to have lower average scores that are compressed for the unknown function category and do not increase dramatically until reaching KFSC >2 true positive status. PFS exhibits higher overall scores compared to S-FPx but shows comparable responsiveness to increasing KFSC. The score distribution of the two models as shown by their inter-quartile ranges also increases by almost 2-fold with increasing KFSC, which is counter to the expectation for increased confidence in classification (**Figure 5C**). Consequently, the recommendation thresholds used for S-PFx and PFS must be low to enable either model to capture even a small percentage of true positive phosphosites. A separate analysis comparing only PFS and SAPH-ire TFx, which includes a larger phosphosite overlap (49,935 phosphosites), showed similar results (**Figure S2**).

In contrast to S-FPx and PFS, the score distributions for SAPH-ire TFx become less, rather than more broad with increasing KFSC, concomitant with the expectation for greater confidence with increasing score (**Figure 5B,C**). ROC and recall curves for all three models show that this difference is largely due to improved recall performance of SAPH-ire TFx, while ROCAUC is otherwise similar between the three different models (**Figure 5D**). The practical consequences of the differences between SAPH-ire TFx and other models is perhaps most evident in terms of the number of missed calls based on recommended thresholds, where as many as 32% of highly confident true positive functional phosphosites (KFSC_MAP_ > 2) are mis-called by previous models – a quantity that is lowered to less than 6% in SAPH-ire TFx (**Figure 5E**). This trend was not specific to whether the phosphosite was a serine, threonine, or tyrosine, further suggesting that SAPH-ire TFx performs equally well regardless of this distinction (**Figures S3**). This also results in an increase in the number of PTMs recommended as functional at all thresholds (**Figure 5F**). Both PFS and SAPH-ire TFx performed equally well for phosphosites whose functionality could have been easily predicted through association with validated functional SLiMs defined by the ELM resource database (**Figure S4**).

We next compared all three models to a recently published fourth model that is not based on machine learning, but rather on a derivative of sequence homology modeling (Strumillo et al., 2018) (**Figure S5**). In brief, this method defines phosphorylation hotspots based on sequence conservation of protein regions in domain families that are densely populated with observed phosphorylation sites. In general, high scores were enriched for conserved phosphosite hotspots regardless of model, with SAPH-ire TFx exhibiting the best overall performance in terms of recall and score distribution across KFSC.

In summary, benchmarking tests of SAPH-ire TFx support the conclusion that the model is robust and effective for the classification of PTM functional status and surpasses the recall performance of previous models.

### Evaluating the model using newly reported experimental evidence and disease linkage

A fortuitous time gap between model development and the writing of this report allowed us to test the accuracy of SAPH-ire TFx predictions using newly reported experimental evidence that arrived after scoring was complete. Between June and December 2019, an update to the functional site database curated by PhosphoSitePlus resulted in an increase of 1066 new functional PTM sites. Consequently, we could use the new data to simulate a situation in which an experimentalist has chosen to investigate the functional impact of a PTM upon recommendations provided by SAPH-ire TFx. In this case, MAPs originally classified as ‘unknown’ in the model output could be re-classified as known functional and then this information used to evaluate model effectiveness. To do this, we cross-referenced the new functional data with existing data from the SAPH-ire TFx expanded dataset, revealing 723 MAP associations (**Figure 6A**). Of these, we further discriminated between two classes: PTMs previously associated with MAPs that were already known to be functional due to association with functionality in other PTMs (Class I; 228) and PTMs associated with MAPs previously unassociated with any functional evidence (Class II; 495). In each class, the curated functional mechanisms regulated by these PTMs were diverse – spanning from regulatory control over molecular association to protein localization, enzyme activity, receptor internalization, and protein degradation/stability (**Figure 6A**).

**Figure 6.**
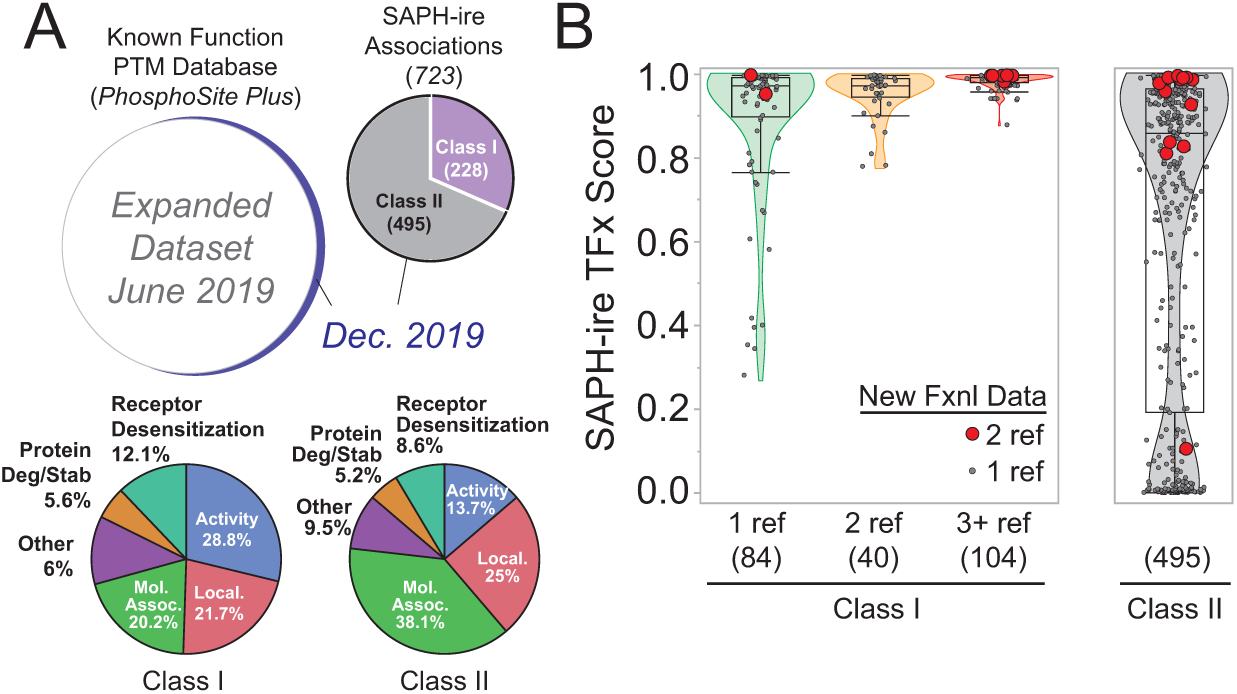
SAPH-ire TFx performance with new functional and disease-linked PTM data. (A, top) Venn diagrams depicting new functional PTM data in comparison to the expanded dataset from figure 2. Class I PTMs are new functional PTM data already associated with known functional MAPs in SAPH-ire TFx. Class II PTMs are new functional PTM data associated with MAPs previously classified as unknown functional (represent completely new experimental data). (A, bottom) pie chart indicating molecular function categories curated for the new functional PTMs. (B) Score distributions for new functional PTM data in Class I and Class II. Red circles correspond to PTMs with 2 references supporting functional impact of the PTM (from the December 2019 update). Original MAP classification (1, 2, 3+ refs) is based on the original classification from figure 2.

The median SAPH-ire TFx score for functional PTMs in class I was above the recommendation threshold for MAPs of known function previously supported by evidence from 1 to 3+ references (**Figure 6B, left**). Moreover, new functional PTMs with more than one reference (from the December 2019 update) were further enriched above the threshold in most cases (red circles). Some of the associated references for the new functional data were from as recent as 2018, which suggest that experimental redundancy within a MAP is common and also probably not always well known to the experimentalist – hence the advantage of tracking function via alignment position in a family. In class II, which represent new functional PTMs that align with MAPs previously classified as unknown function, we found a similar trend (**Figure 6B, right**).

Although the median score was slightly below our lowest threshold of 0.95, the bulk density of the new data scored near or above this threshold. Similarly, most new functional PTM data with more than one reference (red circles) were enriched above the recommendation thresholds for SAPH-ire TFx. Finally, we also noted that several new functional PTMs also scored poorly by SAPH-ire TFx in class II. However, benchmark comparisons against S-PFx and PFS again showed significant improvement in recall of the newly reported functional PTMs by SAPH-ire TFx, suggesting that the model outperforms existing methods (**Figure S6**). Time will be necessary to establish if new reports of PTM function are corroborated by more than one investigation before any further conclusions can be drawn. In summary, new functional PTM data serve as proxies for experimental validation of the SAPH-ire TFx model and provide strong evidence that the model is effective for recommending functional modifications that span a broad range of molecular control mechanisms.

### SAPH-ire TFx in practice: Recommending PTMs of unknown function at the intersection of empirical and computational evidence

Once validated, we decided to use SAPH-ire TFx predictions to filter PTMs that are proximal to functional residues and/or localized within functional short linear motifs (SLiMs). We reasoned that such an effort would highlight PTMs of potentially high biological impact. Therefore, we investigated SAPH-ire TFx-recommended PTMs that are within 3 residues of a pathogenic amino acid substitution mutation (curated by ClinVar) and within a predicted functional SLiM motif (curated by the ELM resource). 517 PTMs within the SAPHire-TFx set met these conditions (**Figure 7A**). Among these, 84 PTMs were already known to be functional and among the remaining 433 unknown function PTMs, 160 (36.9%) were recommended by SAPH-ire TFx (**Figure 7B**; **Table S5**). To assess the biological landscape of the recommended PTMs, we performed a gene ontology (GO) enrichment analysis (http://geneontology.org) of proteins in the recommended list normalized to the GO enrichment of all human PTMs in the extended dataset (Consortium, 2018). Several GO terms from Biological Process, Molecular Function, and Cellular Component GO categories were significantly enriched beyond expectation (up to ∼83-fold after normalization) (**Table S6**). Several major clusters are immediately evident – most notably: muscle, cardiac, heart; morphogenesis & development; as well as transmembrane, receptor, and signaling, among others (**Figure 7C**). This is consistent with our previous observation that several unknown function PTMs are enriched in cardiomyopathies (Torres et al., 2016), and reiterate that this area of biology may be understudied in terms of PTM regulation.

**Figure 7.**
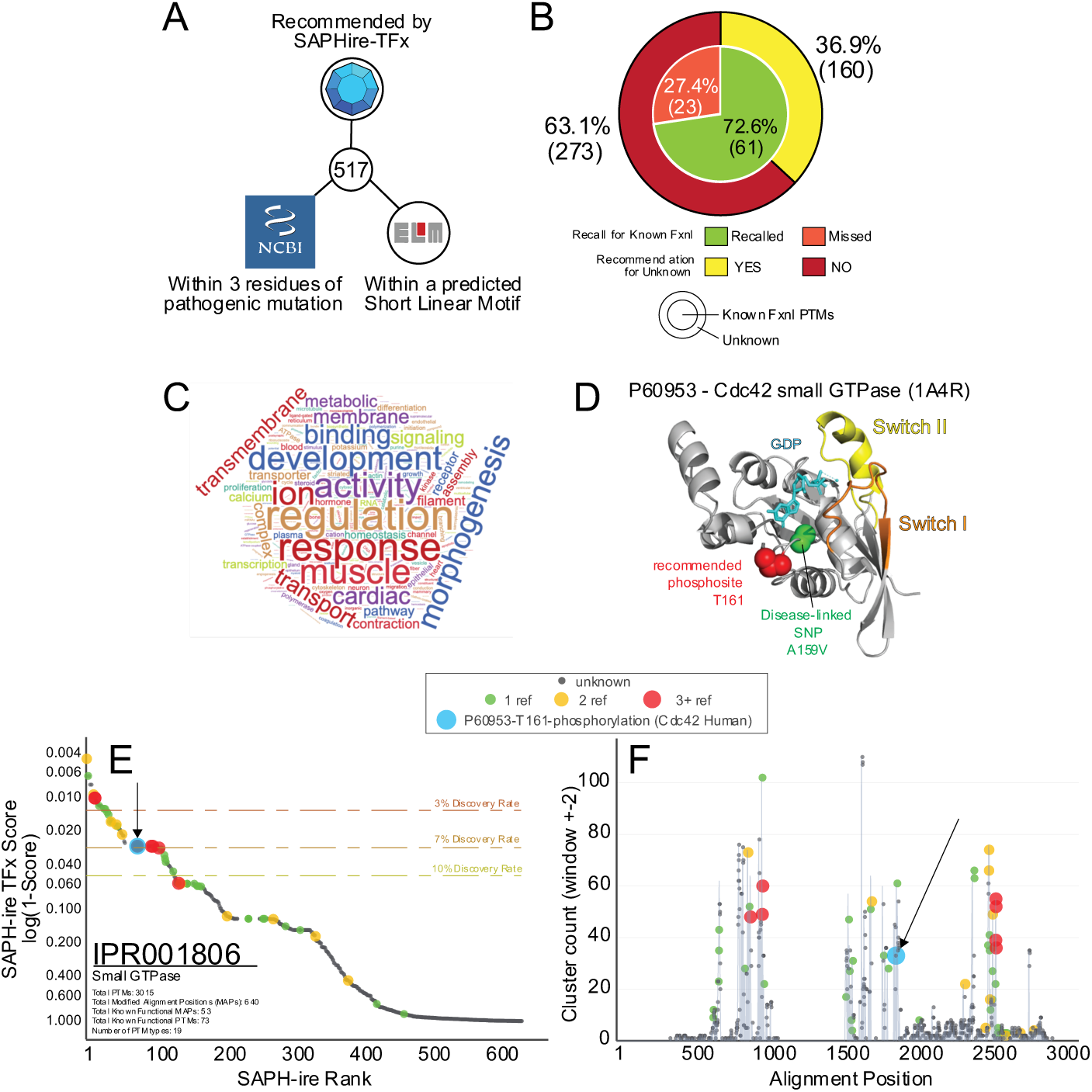
Exploring PTMs at the intersection of multiple independent sources of functional evidence. (A) Schematic diagram depicting the tri-partite filter used for identifying critically important PTMs here. (B) Analysis of recall and recommend rates for the resulting 517 filtered PTMs derived in A. (C) Word cloud diagram showing term frequency within the GO terms enriched between 5x-85x over expectation (greater frequency = larger size). (D) X-ray crystal structure (PDB:1A4R) of human Cdc42 with important regulatory (yellow/orange), PTM (red), and disease-linked mutation sites indicated (green). (E,F) SAPH-ire TFx MAP rank plot and PTM cluster count plot with known and unknown function MAPs indicated by color and circle size (downloaded from https://saphire.biosci.gatech.edu).

Surveying the 160 recommended PTMs of unknown function revealed several hotspots across a wide variety of very important proteins. After filtering further by whether the PTM is in the vicinity of a known functional modification (Known by Neighbor), resulted in 49 distinct PTMs for which we found no evidence of function reported (**Table S5**). Several PTMs in actin and other muscle/heart-related proteins dominated the list. We were particularly surprised to find a phosphosite (T161) in Cdc42, a small GTPase critical for actin dynamics and cell polarity regulation, and an important cancer target (Maldonado and Dharmawardhane, 2018). The recommended site falls very close to the catalytic pocket of the enzyme much like the switch I/II regions that are essential for GTPase activity regulation (**Figure 7D**). We used the SAPH-ire website (https://saphire.biosci.gatech.edu) to view T161 in context of the small GTPase family (IPR001806), finding that the site is one of over 3000 distinct family PTMs and falls within a MAP that ranks in the top 100 (**Figure 7E**). While this site does fall within a small cluster of PTMs in the family, it is one that is understudied compared to other regions of the protein, made obvious by the KFSC markers for known function sources (green, yellow, red circles) (**Figure 7F**). Taken together, these data demonstrate the utility of SAPH-ire TFx as a model and a resource for the study of PTMs in eukaryotes.

## DISCUSSION

We have created a new machine learning model – SAPH-ire TFx – that is capable of confidently recommending PTMs of likely functional significance. The model is shown to be highly predictive for recall of PTMs of known function, and this property is enhanced at increasing recommendation thresholds provided by the model. After its development, we tested the model with an expanded dataset to which it had never been previously exposed, showing that its performance characteristics are robust. To estimate its performance with physiologically relevant predictions, we demonstrated that the model functions adequately to predict the functionality of PTMs curated 6 months after the model was developed and tested – providing a type of meta-experimental validation that goes beyond previously reported models. In a series of benchmarking tests, we further showed that SAPH-ire TFx outperforms existing machine learning or conservation-based hotspot models (including one of our previous models) in all respects, including ROC, recall, and prediction confidence (**Figure 5**). Finally, we provide quantitatively validated thresholds that maximize confidence, recall, and recommendations of unknown function PTMs (the goal of the model).

Through development and validation of SAPH-ire TFx, we have shown that single features often held as the standard for predicting whether or not a PTM is likely to be functional – such as evolutionary conservation or proximity to catalytic residues – are not capable by themselves of capturing the breadth of functional PTM observed over the last several decades. Thus, we suspect that models failing to validate the capture of these true positive data can suffer in their ability to make confident recommendations. By all benchmarking tests conducted, the SAPH-ire TFx model captures the largest swath of known functional PTMs. Evidence from LIME analysis of model feature contributions shows that this is in part due to its ability to capture functional PTMs based on more than one combination of features. Indeed, the model recalls known functional PTMs using either strong evidence from a single feature or weaker evidence across several features (**Figure 3**). Consequently, SAPH-ire TFx exhibits equivalence to other models in the recall of *low hanging fruit*, represented by PTMs whose role in protein function could be easily guessed by conservation, proximity to functional residues or observation frequency (**Figure S3**); however, it significantly outperforms these models in the recall of *high hanging fruit*, represented by PTMs that are not easily recognized as functional by any one single feature alone (**Figure 5, S5**).

Considering its ability to capture a broad range of functional PTM, SAPH-ire TFx shows a considerable increase in the number of PTM sites recommended as likely functional (Figure 4F). Importantly, these recommendations are based on very strict thresholds (score ≥ 0.95, 0.975, 0.985) that capture the top 67% (at most) of all known functional modifications (at score ≥ 0.95) included in the study. If we loosen this threshold to score = 0.75, nearly 90% of currently known functional PTMs are captured. However, even at this loose threshold nearly half (∼47%) of PTMs with unknown function would not be recommended. While we would not conclude that everything below a score of 0.75 is non-functional, we can conclude that PTMs below this loose threshold do not share feature combinations observed for 90% of the functional PTM sites reported thus far. This is striking and suggests multiple possibilities: that a vast majority of studies on the functionality of a PTM have been historically restricted to those falling within a narrow range of specific features (e.g. observation frequency) or that nearly half of all observed PTMs are non-functional *noise* in our biological systems of interest. It’s also possible that SAPH-ire TFx does not efficiently detect PTMs whose function is mediated through interaction with other modifications. We have begun to evaluate the first two hypotheses through experimental validation of SAPH-ire output wherein we empirically test the functionality of PTM sites across the range of SAPH-ire scores regardless of recommendation thresholds (Mukherjee et al., 2019). While our findings have been consistent with the noise hypothesis, more evidence will be necessary to understand this question carefully. Indeed, evidence necessary to train machine learning models to detect combinatorial regulatory modifications is severely limiting. In any case, the SAPH-ire TFx model provides the most comprehensive view of functionality to date.

All of the data described in this report is publicly accessible at https://saphire.biosci.gatech.edu. The site allows investigators to explore several aspects of the SAPH-ire TFx model through customizable graphical or tabular output. This resource includes not only scoring data, but also several other features that are borne from multiple sequence alignment of PTMs (i.e. MAPs) including: the relation to known functional PTM data (neighboring or aligned), protein and family-specific information, PTM type information, density or PTM clustering information, among other outputs that enable one to quickly survey any given protein or protein family for direct and aligned PTM evidence. We have shown an example of the graphical output here (**Figure 7E,F**), and have ensured that capturing these graphics for use by the end user is simple. As a result of these efforts we hope to propel forward the study and understanding of PTM function not only through an improved quantitative model but also through improved accessibility/visualization – both of which are equally important to ensure future progress in the field.

## MATERIALS AND METHODS

### PTMs and multiple sequence alignment

SAPH-ire TFx is PTM agnostic and includes 56 different PTM types (PTMtype), the bulk of which correspond to phosphorylation, ubiquitination, acetylation, methylation, N-linked glycosylation, and sumoylation (**Table S1**). PTMs were collected from multiple sources including PhosphositePlus (Hornbeck et al., 2015), SysPTM (Li et al., 2014), and dbPTM (Huang et al., 2016). Each PTM was mapped to UniProt identifiers (UID) and validated by matching the native position (NP) and residue (res) of the curated PTM to UniProt sequences verified for 100% sequence identity using BLAST (Altschul et al., 1990). Isoforms, although rare in the PTM dataset, were also included. The final PTM dataset for training contained 435,750 unique PTMs (identified by UID-NP-res-PTMtype).

Later this process was repeated with an expanded PTM dataset for the purpose of model validation. UID entries were mapped to whole sequence protein families using InterPro (Mitchell et al., 2015) followed by multiple sequence alignment of family-linked UniProt sequences using MUSCLE with default parameters (Edgar, 2004). Families with fewer than 2 members containing at least 1 PTM per member were excluded. This process resulted in a final 512,015 PTMs mapped to 8,039 families (**Table S2**) containing 38,231 UIDs representing 763 eukaryotic organisms (**Table S3**).

### Feature selection

SAPH-ire features were derived from Modified Alignment Positions (MAPs) corresponding to family alignment positions that harbor at least one PTM, as described previously in detail (Dewhurst and Torres, 2017; Torres, 2016). A total of eleven features were extracted from 319,981 MAPs (containing the 512,015 PTMs) for inclusion in neural network models described below (**Table S4**). The number of unique PTM types observed in the alignment position (AP_PTc), the PTM residue conservation within the alignment position (PRC), the predicted disorder of the modified residue (Dis), and the number of unique modified residues within the alignment position (Modified residue count; MRc) all provide the model with an evolutionary conservation-based perspective on the MAP – a feature that has been shown to be effective in the past (Landry et al., 2009). The next group of features provide information on the local environment of the MAP (not including the MAP in question) by providing the modified residue count of neighboring MAPs (NMrc), the count of neighboring MAPs with modification (Neighbor count, Nc), and neighboring MAPs that harbor known functional PTM (Known neighbor count, KNc). Neighboring residue context has been shown to be an effective predictive feature in the past by us and others (Beltrao et al., 2012; Minguez et al., 2015). Lastly, the number of sources that have reported observation of the PTM (observation source count; OBSrc), a normalized version of this feature that takes family membership into account (OBSrc_pf), and the number of UniProt entries associated with the MAP excluding gaps (Alignment position member count, AP-MEM) are utilized in this model for the first time here.

### Model implementation, cost function optimization, training, and model selection

The SAPH-ire TFx neural network model and modified cost function (defined below) were implemented in Tensorflow using the estimator API (Abadi et al., 2016). PTMs and MAPs were processed into features using Python 3.7.3 and Pandas 0.24.0 (McKinney, 2010). MAPs with at least two references (PMIDs) of corroborating evidence of biological function, defined by PhosphositePlus (Hornbeck et al., 2015), were treated as the positive class with all others being treated as negative. MAPs with a single source were treated as negative because they lacked independent confirmation of functional significance and because their inclusion weakened model performance. The cost for false negatives (misclassified known functional PTMs) was weighted at a 4 to 1 ratio to false positives (PTMs with unknown function classified as functional) to reflect the goal of recommending unstudied PTMs for research. Below this 4:1 ratio the performance suffered, and above it there was no significant improvement while the model began to exhibit signs of overfitting.

Neural network models of various structure (in terms of connectivity, activation function, cost function weighting, etc.) were generated in batches of 100 or 200 depending on architecture complexity. The training set for these models were bootstrapped in order to over-represent the positive class to avoid sample distribution biasing (Dupret and Koda, 2001). Models were trained using a 33% holdback rate, with this holdback being used to evaluate batches of models against the same evaluation set. Model selection relied on Receiver Operating Characteristic (ROC) and Recall summarization metrics integrated by an Fzero score defined as:

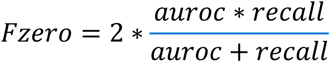

Optimal models were defined as those with the greatest Fzero score.

### SAPH-ire TFx model architecture and optimizing performance through recall

For model training, we used a SAPH-ire training dataset generated in 2018, consisting of 435,750 PTMs coalesced by multiple sequence alignment into 272,968 MAPs (**Figure 8A**). From each of more than a dozen architecture and training permutations, 200 models were stochastically trained and evaluated by Fzero and Recall and then filtered to identify the most consistently high performing architectures (Batch 1). Due to the intended goal of SAPH-ire to identify “potential positives”, a high precision (true positives captured / total positives) was not only unwanted but indicative of a poor model and therefore not included as a summary metric. The agreement between most of the models made collective intelligence approaches redundant. Therefore, an additional series of models trained with an added feature (family-weighted observation source count) were generated (Batch 2). In general, the top model from each batch varied only slightly in terms of ROC AUC (0.68 – 0.73), but varied dramatically in recall (0.5 – 0.8), suggesting that significant gains in model performance were achieved by considering recall in addition to ROC AUC (**Figure 8B**). The top performing models from these two independent evaluations were averaged together to represent the final SAPH-ire score, which outperformed either top model alone (**Figure 8B** (inset)). This final score resulted in excellent predictive (AUC = 0.792) and recall (AUC = 0.694) performance (**Figure 8C**). The top model from Batch 1 (Model 1; representing a global perspective) and Batch 2 (Model 2; representing a local perspective) rely on the same architecture in which the input layer flows into two hidden layers consisting of a rectified linear unit (RELU) followed by a saturating tanh function (**Figure 8D**). The RELU allows for scaling the inputs and dampening the impact of large differences in the magnitude of the features, while the tanh layer compresses the output to a fixed probability distribution. Model 1 takes in 10 features without pre-processing, allowing it to use the network architecture to scale the inputs globally. In contrast, Model 2 uses 11 features, with observation source count normalized by family for each MAP serving as the additional feature and the other 10 features normalized globally. The output of both models are averaged together to give the SAPH-ire TFx score.

**Figure 8.**
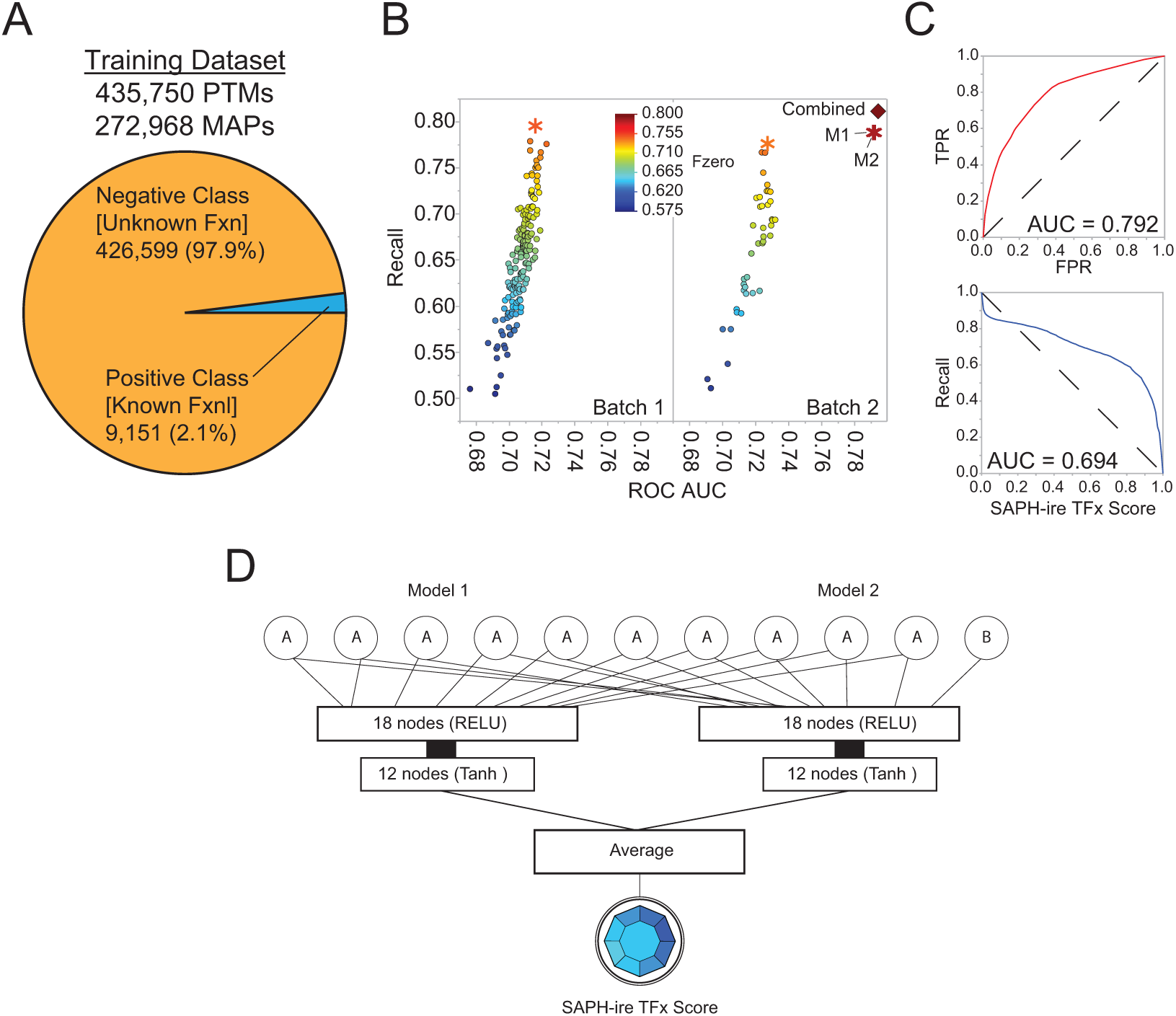
Numerical summary of SAPH-ire TFx development, training, and performance. (A) Diagram of the training dataset with positive and negative classes indicated. Note that Modified Alignment Positions (MAPs) with less than 2 literature sources of support were included in the negative class as they were not used at any point for training the model (see methods). (B) Plot of true positive recall versus ROC AUC versus Fzero (color) for the top two of 200 models generated using ten (Batch 1) or eleven (Batch 2) features (MAP-level analysis). Dashed box (on same scale) indicates performance of individual top models from Batch 1 and 2 (M1, M2) and a combined model (SAPH-ire TFx) that is generated by taking the average score of M1 and M2 on the expanded dataset (shown further below). (C) ROC and recall threshold curves for the combined model (based on MAP-level analysis). (D) Final model architecture, in which the 10 shared input features [A] and 1 unique input feature [B] of M1 and M2 are indicated.

### Pathogenic SNP and Motif Enrichment Analysis

Genetic mutations and the curated interpretation of their significance to disease were collected from Clinvar (Landrum et al., 2018). The proteins affected by genetic mutations were filtered for single nucleotide polymorphisms (SNPs) that alter PTM sites present within the SAPH-ire dataset. These sites were then aggregated by SAPH-ire MAP and separated into one of four Clinvar-designated categories: Benign, Likely Benign, Pathogenic, or Likely Pathogenic. Only pathogenic categories were used for further analysis.

Experimentally validated Short Linear Motifs (SliMs) were collected from The Eukaryotic Linear Motif (ELM) resource for functional sites in proteins (Gouw et al., 2018). At the time of this study, ELM contained 289 motif classes clustered into 6 motif categories based on functional assessment of 3,523 validated instances. Only PTMs that occurred within a validated motif instance in any category were used for the motif enrichment analysis.

In order to identify additional instances of motifs outside of the experimentally validated set provided by ELM, we scanned the proteins contributing to the SAPH-ire TFx dataset for amino acid sequences matching the regular expression patterns provided by ELM. As a purely regular expression based approach would produce numerous false positives, the resulting SLiMs were filtered based on conservation of the detected motif within a multiple sequence alignment of the protein family, only keeping those that had more than 40% occurrence at precise alignment positions within the family – as prescribed by ELM curators previously (Gibson et al., 2015).

To investigate PTMs at the intersection of functional SLiM motifs, pathogenic SNP mutations and SAPHire-TFx recommendations, PTMs within the expanded dataset were filtered based on the following criteria: (1) The PTM must be recommended by SAPHire-TFx; (2) The PTM must be within 3 residues of a pathogenic SNP mutation that changed the amino acid sequence of the associated protein; and (3) The PTM must reside within a predicted SLiM that has passed the previously stated regular expression filters.

### Graphical and statistical data analyses

Graphical and statistical data analyses were achieved using a combination of R (R Core Team, 2013), Python (specifically the pandas library) (McKinney, 2010), and JMP 14.1 (SAS Institute Inc.).

### SAPH-ire Website

The SAPH-ire website (https://saphire.biosci.gatech.edu) is composed of three microservices managed by Docker (https://www.docker.com/community/open-source). The SAPH-ire dataset including predictions is loaded into a MongoDB microservice (https://www.mongodb.com/), which is then queried dynamically by Gunicorn microservice (https://github.com/benoitc/gunicorn). The Gunicorn microservice serves as the API which is accessible directly at the api endpoint of the SAPH-ire site with structured queries. The API is read by a visualization microservice developed using Vue.js (You, n.d.), Plotly (Inc., 2015), and Vuetify (Leider, 2020). Vue.js was used to create the interactive single page application, Vuetify provided reactive application components, and Plotly provided dynamic graph element.

## ACKNOWLEDGEMENTS

We would like to thank Zahra Nassiri Toosi, Wei Li, and other members of the Torres lab for careful review and beta-testing of the SAPH-ire website. Special thanks also to Jiani Long and Ragy Haddad for pilot work on the model and website conceptualization. Special thanks to Dr. Peng Qiu of Georgia Institute of Technology for critical review of the manuscript and contributions to model design. This work was funded by National Institutes of Health R01 GM117400 and to M.T.

## SUPPLEMENTAL TABLES AND FIGURES

*Supplemental tables can be found as individual tabs in the supplemental excel file*.

**Table S1**. Frequency of PTM types analyzed by SAPH-ire TFx.

**Table S2**. List of InterPro families analyzed in SAPH-ire TFx.

**Table S3**. List of organisms represented by SAPH-ire TFx.

**Table S4**. Description of features used in the SAPH-ire TFx model.

**Table S5**. List of PTMs that intersect between SAPH-ire TFx, ELM, and Clinvar.

**Table S6**. GO enrichment analysis of 221 TFx-recommended PTMs from Table S5.

**Figure S1.**
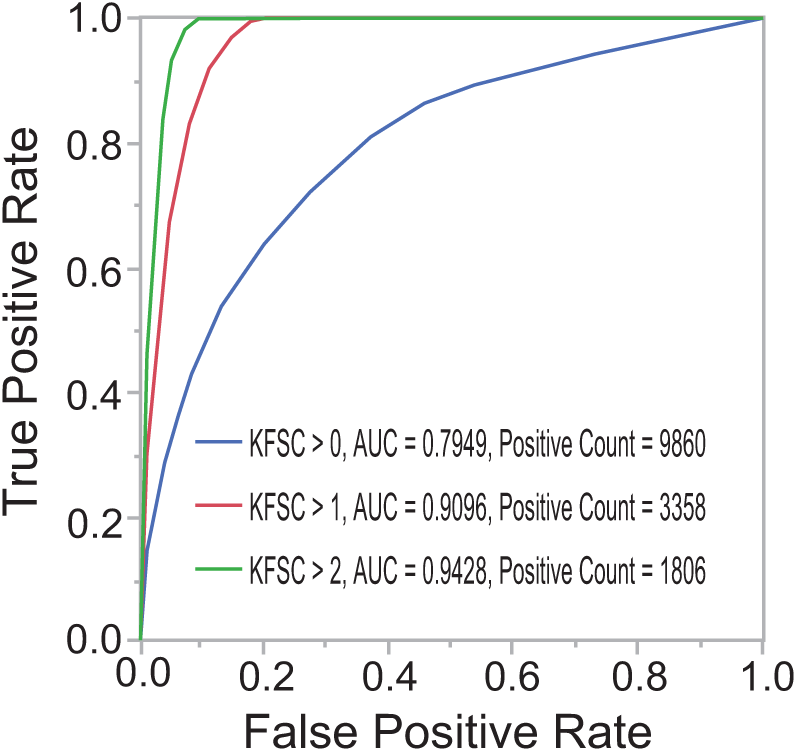
ROC curves at different KFSC thresholds. (unfurled in Figure 1E).

**Figure S2.**
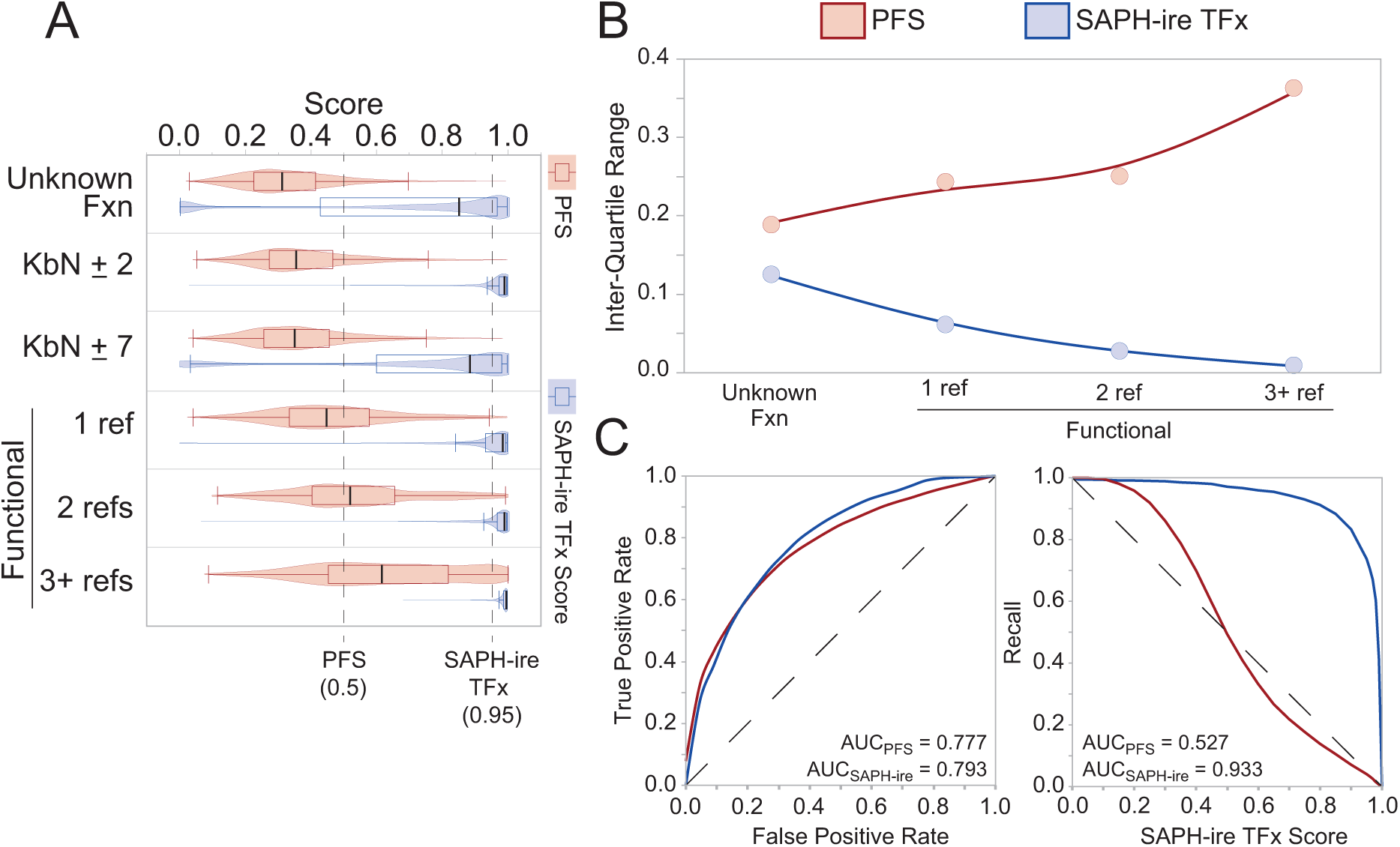
Pairwise comparison between SAPH-ire TFx and the PFS machine learning models. These data include 49,935 phosphosites that overlap between the SAPH-ire TFx and PFS datasets. (A) Score distribution relative to functional status. (B) Inter-quartile ranges from the distributions in A. (C). ROC and recall curves for the score comparison of each model.

**Figure S3.**
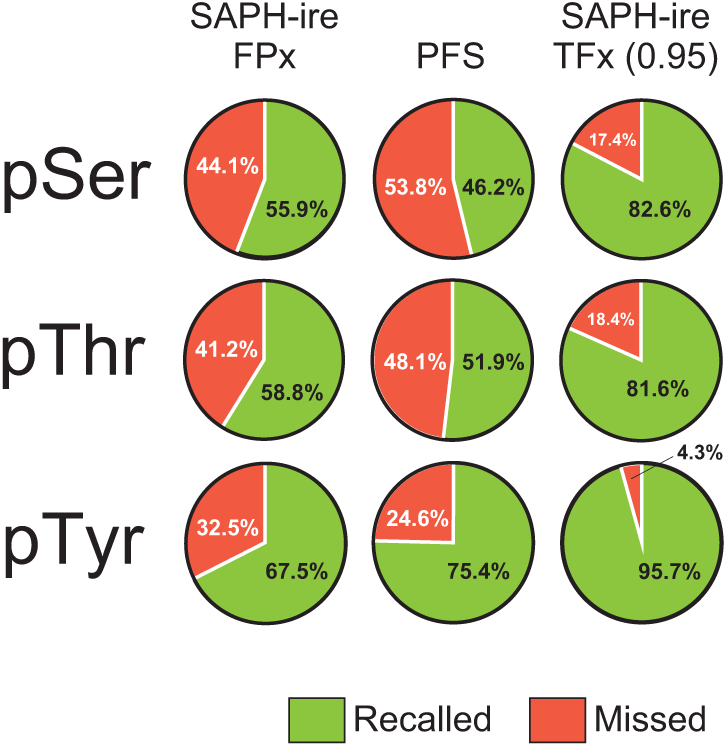
Recall performance improvements observed with SAPH-ire TFx are independent of whether the site is serine, threonine or tyrosine. A total of 24,695 phosphosites that overlap between S-PFx, PFS, and SAPH-ire TFx were parsed by site identity and the percent recalled or missed tallied based on model-specific thresholds indicated in Figure 4.

**Figure S4.**
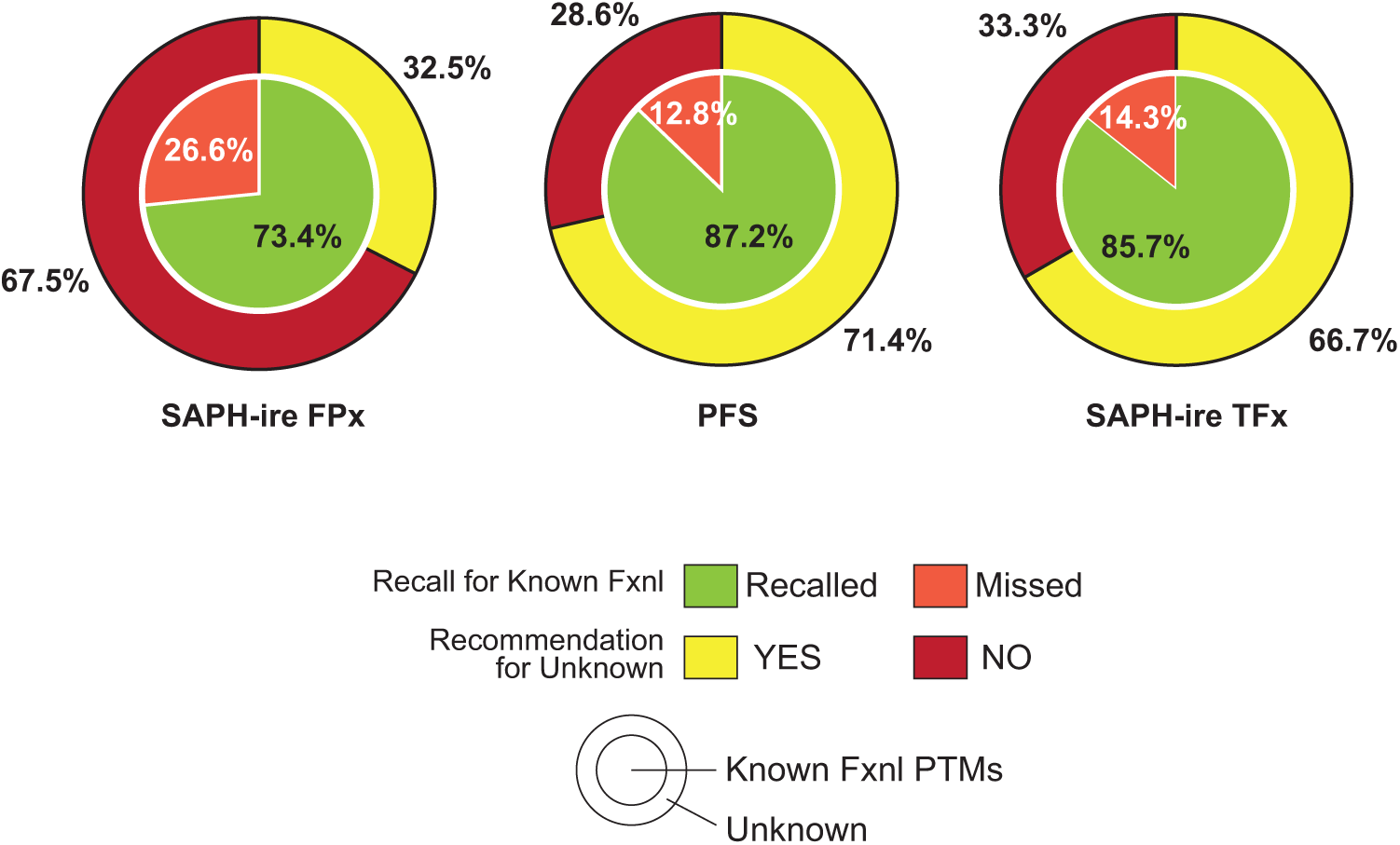
Model recall and recommendation comparison for PTMs associated with validated functional SLiMs from the ELM resource.

**Figure S5.**
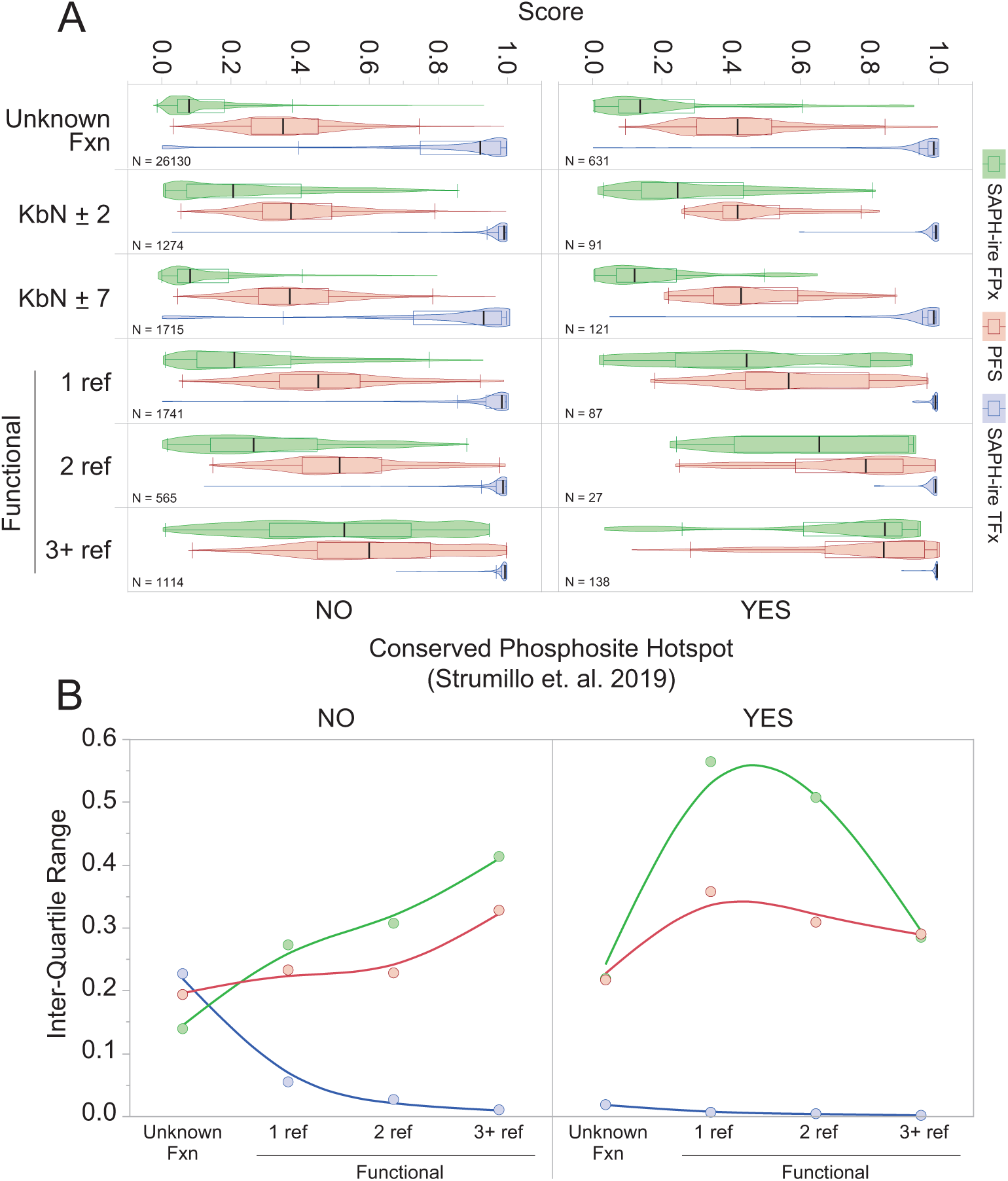
Comparison of SAPH-ire FPx, PFS, and SAPH-ire TFx models relative to phosphosite conservation hotspot analysis. Phosphosites localized within conserve phosphosite hotspots predicted by Strumillo et al. were used to bin data from the three-model comparison shown in figure 5. (A) Score distribution relative to functional status relative to predicted hotspots. (B) Inter-quartile ranges from the distributions in A.

**Figure S6.**
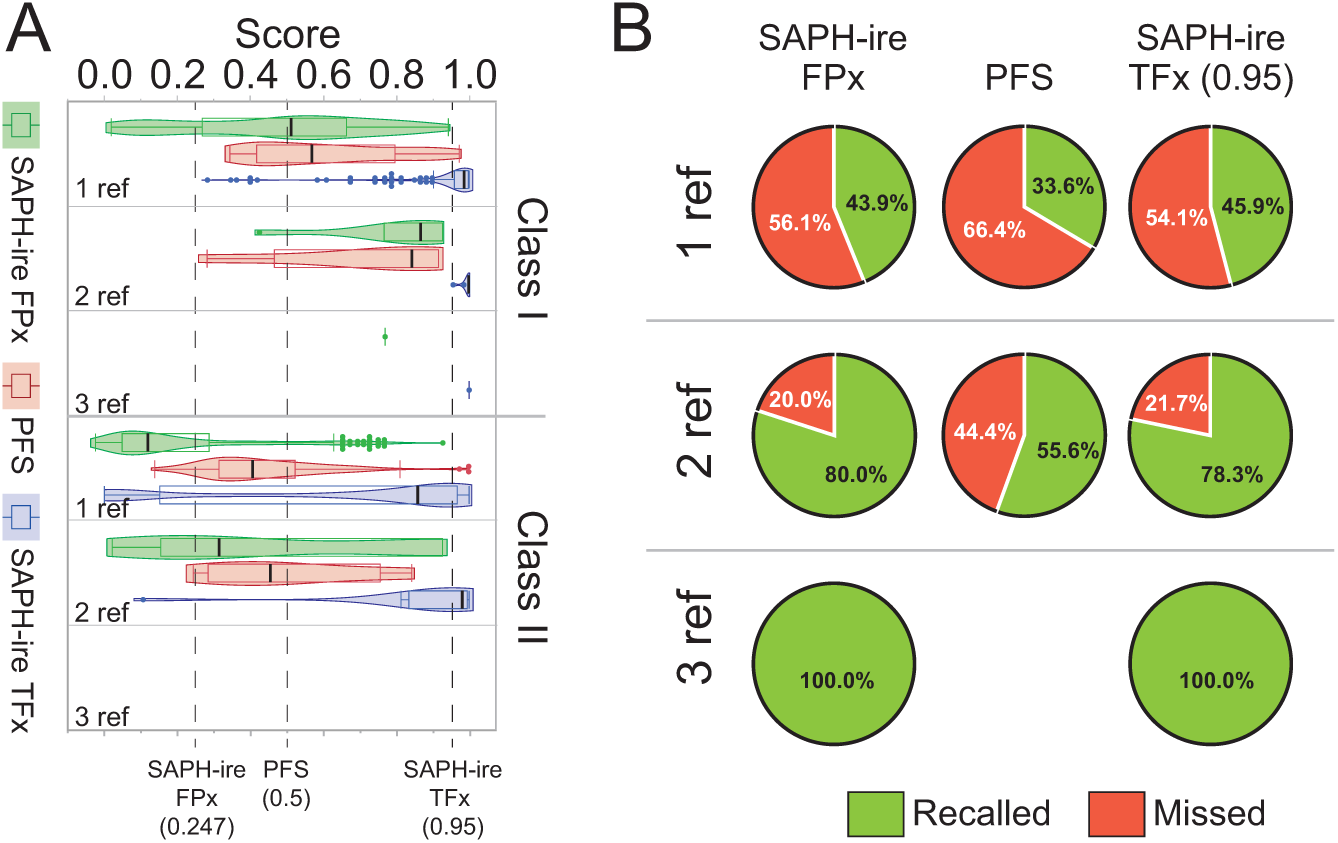
Model comparison for Class I and II newly curated functional phosphosites. (A) Model score distributions for Class I and II newly curated functional PTMs (see figure 4A). “1, 2, 3 refs” refers to number of PMIDs associated with the new data only (not the data from the original analysis of the extended dataset). (B) Recall rates for each model is shown relative to number of references supporting functionality of the PTM from A (KFSC = 1, 2, 3 ref).

## REFERENCES

Abadi M, Barham P, Chen J, Chen Z, Davis A, Dean J, Devin M, Ghemawat S, Irving G, Isard M, Kudlur M, Levenberg J, Monga R, Moore S, Murray DG, Steiner B, Tucker P, Vasudevan V, Warden P, Wicke M, Yu Y, Zheng X. 2016. TensorFlow: A system for large-scale machine learning.

Altschul SFF, Gish W, Miller W, Myers EWW, Lipman DJJ. 1990. Basic local alignment search tool. J Mol Biol 215:403–10. doi:10.1016/S0022-2836(05)80360-2

Beltrao P, Albanèse V, Kenner LR, Swaney DL, Burlingame A, Villén J, Lim W a, Fraser JS, Frydman J, Krogan NJ. 2012. Systematic functional prioritization of protein posttranslational modifications. Cell 150:413–25. doi:10.1016/j.cell.2012.05.036

Chen C, Huang H, Wu CH. 2017. Protein Bioinformatics Databases and Resources. Methods Mol Biol 1558:3–39. doi:10.1007/978-1-4939-6783-4_1

Consortium TGO. 2018. The Gene Ontology Resource: 20 years and still GOing strong. Nucleic Acids Res 47:D330–D338. doi:10.1093/nar/gky1055

Csizmok V, Forman-Kay JD. 2018. Complex regulatory mechanisms mediated by the interplay of multiple post-translational modifications. Curr Opin Struct Biol 48:58–67. doi:10.1016/j.sbi.2017.10.013

Dewhurst HM, Choudhury S, Torres MP. 2015. Structural Analysis of PTM Hotspots (SAPH-ire)--A Quantitative Informatics Method Enabling the Discovery of Novel Regulatory Elements in Protein Families. Mol Cell Proteomics 14:2285–97.

Dewhurst HM, Torres MP. 2017. Systematic analysis of non-structural protein features for the prediction of PTM function potential by artificial neural networks. PLoS One 12:e0172572. doi:10.1371/journal.pone.0172572

Dupret G, Koda M. 2001. Bootstrap re-sampling for unbalanced data in supervised learning. Eur J Oper Res 134:141–156. doi:10.1016/S0377-2217(00)00244-7

Edgar RC. 2004. MUSCLE: multiple sequence alignment with high accuracy and high throughput. Nucleic Acids Res 32:1792–1797. doi:10.1093/nar/gkh340

Gibson DG, Glass JI, Lartigue C, Noskov VN, Chuang R-Y, Algire M a, Benders G a, Montague MG, Ma L, Moodie MM, Merryman C, Vashee S, Krishnakumar R, Assad-Garcia N, Andrews-Pfannkoch C, Denisova E a, Young L, Qi Z-Q, Segall-Shapiro TH, Calvey CH, Parmar PP, Hutchison C a, Smith HO, Venter JC. 2010. Creation of a bacterial cell controlled by a chemically synthesized genome. Science 329:52–6. doi:10.1126/science.1190719

Gibson TJ, Dinkel H, Van Roey K, Diella F. 2015. Experimental detection of short regulatory motifs in eukaryotic proteins: tips for good practice as well as for bad. Cell Commun Signal 13:42. doi:10.1186/s12964-015-0121-y

Gouw M, Michael S, Sámano-Sánchez H, Kumar M, Zeke A, Lang B, Bely B, Chemes LB, Davey NE, Deng Z, Diella F, Gürth C-M, Huber A-K, Kleinsorg S, Schlegel LS, Palopoli N, Roey K V, Altenberg B, Reményi A, Dinkel H, Gibson TJ. 2018. The eukaryotic linear motif resource – 2018 update. Nucleic Acids Res 46:D428–D434. doi:10.1093/nar/gkx1077

Hornbeck P V, Zhang B, Murray B, Kornhauser JM, Latham V, Skrzypek E. 2015. PhosphoSitePlus, 2014: mutations, PTMs and recalibrations. Nucleic Acids Res 43:D512–20. doi:10.1093/nar/gku1267

Huang K-Y, Su M-G, Kao H-J, Hsieh Y-C, Jhong J-H, Cheng K-H, Huang H-D, Lee T-Y. 2016. dbPTM 2016: 10-year anniversary of a resource for post-translational modification of proteins. Nucleic Acids Res 44:D435-D446. doi:10.1093/nar/gkv1240

Inc. PT. 2015. Collaborative data science.

Johnson JR, Santos SD, Johnson T, Pieper U, Strumillo M, Wagih O, Sali A, Krogan NJ, Beltrao P. 2015. Prediction of Functionally Important Phospho-Regulatory Events in Xenopus laevis Oocytes. PLOS Comput Biol 11:e1004362. doi:10.1371/journal.pcbi.1004362

Landrum MJ, Lee JM, Benson M, Brown GR, Chao C, Chitipiralla S, Gu B, Hart J, Hoffman D, Jang W, Karapetyan K, Katz K, Liu C, Maddipatla Z, Malheiro A, McDaniel K, Ovetsky M, Riley G, Zhou G, Holmes JB, Kattman BL, Maglott DR. 2018. ClinVar: improving access to variant interpretations and supporting evidence. Nucleic Acids Res 46:D1062-D1067. doi:10.1093/nar/gkx1153

Landry CR, Levy ED, Michnick SW. 2009. Weak functional constraints on phosphoproteomes. Trends Genet 25:193–7. doi:10.1016/j.tig.2009.03.003 Leider J. 2020. Vuetify.

Li J, Jia J, Li H, Yu J, Sun H, He Y, Lv D, Yang X, Glocker MO, Ma L, Yang J, Li L, Li W, Zhang G, Liu Q, Li Y, Xie L. 2014. SysPTM 2.0: an updated systematic resource for post-translational modification. Database (Oxford) 2014:bau025. doi:10.1093/database/bau025

Maldonado M del M, Dharmawardhane S. 2018. Targeting Rac and Cdc42 GTPases in Cancer. Cancer Res 78:3101–3111. doi:10.1158/0008-5472.CAN-18-0619

McKinney W. 2010. Data Structures for Statistical Computing in Python.

Minguez P, Letunic I, Parca L, Bork P. 2013. PTMcode: a database of known and predicted functional associations between post-translational modifications in proteins. Nucleic Acids Res 41:D306–11. doi:10.1093/nar/gks1230

Minguez P, Letunic I, Parca L, Garcia-Alonso L, Dopazo J, Huerta-Cepas J, Bork P. 2015. PTMcode v2: a resource for functional associations of post-translational modifications within and between proteins. Nucleic Acids Res 43:D494-502. doi:10.1093/nar/gku1081

Minguez P, Parca L, Diella F, Mende DR, Kumar R, Helmer-Citterich M, Gavin A-C, van Noort V, Bork P. 2012. Deciphering a global network of functionally associated post-translational modifications. Mol Syst Biol 8:599. doi:10.1038/msb.2012.31

Mitchell A, Chang H, Daugherty L, Fraser M, Hunter S, Lopez R, Mcanulla C, Mcmenamin C, Nuka G, Pesseat S, Sangrador-vegas A, Scheremetjew M, Rato C, Yong S, Bateman A, Punta M, Attwood TK, Sigrist CJ a., Redaschi N, Rivoire C, Xenarios I, Kahn D, Guyot D, Bork P, Letunic I, Gough J, Oates M, Haft D, Huang H, Natale D a., Wu CH, Orengo C, Sillitoe I, Mi H, Thomas PD, Finn RD. 2015. The InterPro protein families database : the classification resource after 15 years. Nucleic Acids Res 43:D213–D221. doi:10.1093/nar/gku1243

Mukherjee K, English N, Meers C, Kim H, Jonke A, Storici F, Torres M. 2019. Systematic analysis of linker histone PTM hotspots reveals phosphorylation sites that modulate homologous recombination and DSB repair. DNA Repair (Amst) 86:102763. doi:10.1016/j.dnarep.2019.102763

Ochoa D, Jarnuczak AF, Gehre M, Soucheray M, Kleefeldt AA, Vieitez C, Hill A, Garcia-Alonso L, Swaney DL, Vizcaino JAA, Noh K-M, Beltrao P. 2019. The functional landscape of the human phosphoproteome. bioRxiv 541656. doi:10.1101/541656

Ochoa D, Jarnuczak AF, Viéitez C, Gehre M, Soucheray M, Mateus A, Kleefeldt AA, Hill A, Garcia-Alonso L, Stein F, Krogan NJ, Savitski MM, Swaney DL, Vizcaíno JA, Noh K-M, Beltrao P. 2020. The functional landscape of the human phosphoproteome. Nat Biotechnol 38:365–373. doi:10.1038/s41587-019-0344-3

Pascovici D, Wu JX, McKay MJ, Joseph C, Noor Z, Kamath K, Wu Y, Ranganathan S, Gupta V, Mirzaei M. 2018. Clinically Relevant Post-Translational Modification Analyses-Maturing Workflows and Bioinformatics Tools. Int J Mol Sci 20. doi:10.3390/ijms20010016

Prabakaran S, Lippens G, Steen H, Gunawardena J. 2012. Post-translational modification: Nature’s escape from genetic imprisonment and the basis for dynamic information encoding. Wiley Interdiscip Rev Syst Biol Med. doi:10.1002/wsbm.1185

R Core Team. 2013. R: A language and environment for statistical computing.

Reimand J, Bader GD. 2014. Systematic analysis of somatic mutations in phosphorylation signaling predicts novel cancer drivers. Mol Syst Biol 9:637–637. doi:10.1038/msb.2012.68

Reimand J, Wagih O, Bader GD. 2015. Evolutionary constraint and disease associations of post-translational modification sites in human genomes. PLoS Genet 11:e1004919. doi:10.1371/journal.pgen.1004919

Strumillo MJ, Oplova M, Vieitez C, Ochoa D, Shahraz M, Busby BP, Sopko R, Studer RA, Perrimon N, Panse VG, Beltrao P. 2018. Conserved phosphorylation hotspots in eukaryotic protein domain families. bioRxiv 391185. doi:10.1101/391185

Strumillo MJ, Oplová M, Viéitez C, Ochoa D, Shahraz M, Busby BP, Sopko R, Studer RA, Perrimon N, Panse VG, Beltrao P. 2019. Conserved phosphorylation hotspots in eukaryotic protein domain families. Nat Commun 10:1977. doi:10.1038/s41467-019-09952-x

Torres M. 2016. Chapter Two - Heterotrimeric G Protein Ubiquitination as a Regulator of G Protein SignalingProgress in Molecular Biology and Translational Science. pp. 57–83. doi:10.1016/bs.pmbts.2016.03.001

Torres MP, Dewhurst H, Sundararaman N. 2016. Proteome-wide Structural Analysis of PTM Hotspots Reveals Regulatory Elements Predicted to Impact Biological Function and Disease. Mol Cell Proteomics 15:3513–3528. doi:10.1074/mcp.M116.062331

Xiao Q, Miao B, Bi J, Wang Z, Li Y. 2016. Prioritizing functional phosphorylation sites based on multiple feature integration. Sci Rep 6:24735. doi:10.1038/srep24735

You E. n.d. VueJs.

